# Comparative transcriptome and proteome profiling of the chlorophyll metabolism pathway in four cultivars of *Magnolia officinalis*

**DOI:** 10.1101/470880

**Authors:** Qinahua Wu, Dan Wei, Yuping Liu, Chaoxiang Ren, Qianqian Liu, Cuiping Chen, Jiang chen, Jin Pei

## Abstract

*Magnolia officinalis* is an important herb in Chinese medicine that has good therapeutic effects on gastrointestinal motility and helps regulate the spleen and stomach. *Magnolia officinalis* Rehd.et Wils. (abbreviated as CH)、 *Magnolia officinalis* Rehd.et Wils. var. biloba’ DaAoye’ (abbreviated as DA)、 *Magnolia officinalis* Rehd.et Wils. var. biloba‘XiaoAoye’ (abbreviated as XA)、*Magnolia officinalis* Rehd.et Wils. var. biloba‘Liuye’ (abbreviated as LY) are the four main cultivars in Dujiangyan city, Sichuan province in China, and DA has better medicinal effects and higher levels of green pigment than the other three cultivars. Proteomics research not only provides new insights into protein expression patterns but also enables the identification of many attractive candidates for further research on the effects of pesticides on chlorophyll metabolism in *Magnolia officinalis*. Transcriptome and proteome analysis of the four main *Magnolia officinalis* cultivars in Sichuan province in China identified 125,100 unigenes and 6,012 proteins. Proteomic data and parallel reaction monitoring (PRM) revealed that porphobilinogen deaminase, uroporphyrinogen decarboxylase, oxygen-dependent coproporphyrinogen-III oxidase, protoporphyrinogen oxidase 1, magnesium protoporphyrin IX methyltransferase, magnesium-protoporphyrin IX monomethyl ester oxidative cyclase, and protochlorophyllide reductase-like protein, all of which are involved in chlorophyll synthesis, have higher expression levels in DA than in the other three cultivars. This finding is consistent with the observation that DA has the highest concentration of chlorophyll (37.56 mg/g Fw) of the four cultivars. This research is beneficial to the understanding of the differences in the growth of the four cultivars on the molecular level.

## Introduction

*Magnolia officinalis* is a unique and precious tree species in China. It is also the source of a traditional Chinese medicinal herb widely used for almost 2000 years to regulate gastrointestinal motility as well as the functioning of the spleen and stomach. Its barks is the main raw material for a number of remedies used in traditional Chinese medicine, such as Patchouli zhengqi oral liquid, Chaihu shugan pill and stomachic pill with cyperus and amomumet. In addition, it was also widely used in Japan, Korea and southeast Asia.

Leaves are the key organs for plant survival and growth, and their traits reflect the self-regulatory capacity of plants to adapt to complex environmental conditions[1]. Many metabolic products, such as magnolol and honokiol, are able to transfer into bark and underlie the functioning of traditional medicines. However, the regulation of raw material used in medicines that are present in leaves remain unclear. Interestingly, the four main cultivars of *Magnolia officinalis* (abbreviated as CH/DA/XA and LY)in Sichuan province of China have different colours, and previous research has shown that medicine derived from each of these cultivars show significant differences in their action. The lack of genomic data for *Magnolia officinalis* has led to difficulties in the research into the different molecular mechanism of chlorophyll metabolism in each of the four main cultivars found in Sichuan province in China.

In this study, a transcriptome database was constructed, which resulted in the identification of 125,100 unigenes. Subsequently, an LC-MS/MS-based shot-gun proteomics approach was used to quantify the differences in proteins levels in leaves from the four cultivars based on the *Magnolia officinalis* transcriptome database. Proteomic analysis identified 6,012 proteins involved in mechanisms related to chlorophyll and compared their levels in DA and the three other cultivars. Leaf colour is the comprehensive result of the mixing of different pigments. The ability of a plant to synthesize chlorophyll plays an important role in the chlorophyll content of the plant [2]. In the chlorophyll synthesis pathway, Hem C play an important role in the synthesis of the chlorophyll precursor, and subsequently, Hem E （Coproporphyrinogen oxidase (EC 1.3.3.3)）is involved in the sixth step of Heme biosynthesis [3]. In the chlorophyll synthesis pathway, Hem Y (protoporphyrinogen IX) serve to catalyse the oxidation of protoporphyrinogen IX to protoporphyrin IX in yeast cells [4]. Proteomic research of DA, XA, CH and LY revealed that the amounts of Hem C, Hem E and Hem Y show clear differences when compared between DA and the other three cultivars (LY, CH, and XA). The green colour of the leaves of a plant is caused by high levels of chlorophyll and low levels of other pigments. Chlorophyll synthase plays an important role in regulating the whole chlorophyll synthetic process, including chlorophyll binding proteins[5]. Due to the high level of green colour in the leaves of DA, which has leaves that are of a much deeper green than the other three cultivars, porphobilinogen deaminase, uroporphyrinogen decarboxylase, oxygen-dependent coproporphyrinogen-III oxidase, protoporphyrinogen oxidase 1, magnesium protoporphyrin IX methyltransferase (m.58550), magnesium-protoporphyrin IX monomethyl ester oxidative cyclase, and protochlorophyllide reductase-like, all of which are involved in the synthesis of chlorophyll, were quantified and re-identified based on parallel reaction monitoring (PRM) analysis.

The analysis of crosstalk between the transcriptome and proteome of each of the four *Magnolia officinalis* cultivars revealed the processes underlying the molecular regulation of chlorophyll metabolism in leaves and indicated the presence of a relationship between a high level of green colour in leaves and faster growth in *Magnolia officinalis* cultivars. This research is beneficial to the understanding of the differences in the growth of the four cultivars on the molecular level.

## Materials and Methods

### Experimental Materials

The four main *Magnolia officinalis* cultivars (CH, DA, XD and LY) were picked from Dujiangyan city, Sichuan province in China. Three leaves from three plants of each cultivar, and leaves were from the same age, developmental stage that were also in the same leaf position on the tree were collected, quick-frozen in liquid nitrogen, and kept at −80°C after storage for later use. Three leaves of each cultivars were mixed uniformly as a sample for protein and transcriptome analysis.

### Determination of leaf color and chlorophyll content

The amount of chlorophyll in the leaves from DA, XA, LY, and CH cultivars was determined using a 730N-type spectrophotometer according to the method described by [6] Three biological replicates were analysed, and the chlorophyll a, chlorophyll b and total chlorophyll contents were determined and expressed as mg/g FW.

### Library preparation and sequencing

Total RNA was extracted from the leaves using TRizol reagent (Invitrogen, CA, USA). To remove DNA, an aliquot of total RNA was treated with DNase (Takara, Dalian, China). The same amount of RNA from each sample was mixed for sequencing analysis. To ensure the accuracy of the sequencing data, a Nanodrop was used to measure the RNA purity (OD 260/280), concentration, and nucleic acid absorption peak, and an Agilent 2100 was used to determine the RNA integrity using detection indicators that included an RIN value of 28S/18S.

### RNA isolation and transcriptome analysis

Total RNA was extracted from the LM of each of the four cultivars of *Magnolia officinalis* using a total RNA extraction kit (QIAGEN, 74903/4) according to the manufacturer’s instructions. The RNA quantity and integrity were verified using a NanoDrop 2000 spectrophotometer and Agilent 2100 bioanalyzer (Agilent Technologies, Santa Clara, CA, USA). A TruSeq RNA Sample Preparation kit v2 (Illumina, San Diego, CA, USA) was used to construct the cDNA libraries. Subsequently, the libraries were sequenced using an Illumina HiSeq instrument (Illumina, San Diego, CA, USA) that generated 100 bp paired-end reads.

### De novo assembling and functional annotation of reads

The raw sequence reads obtained for each sample were trimmed and assembled de novo using Trinity, and then Tgicl was used to cluster the transcripts into unigenes. To annotate the NT, NR, KOG, and KEGG genes obtained from the transcriptomic analysis, Blastn, Blastx or Diamond was used to align the unigenes based on the SwissProt database. Blast2 GO with NR annotation was used for the GO annotation, while InterProScan5 was used for the InterPro annotation.

### Protein preparations

The samples were first ground in liquid nitrogen, and then the resulting cell powder was transferred to a 5 mL centrifuge tube containing lysis buffer (8 M urea, 2 mM EDTA, 10 mM DTT and 1% protease inhibitor cocktail) and sonicated three times on ice using a high intensity ultrasonic processor (Scientz). The remaining debris was removed by centrifugation at 20,000 × g for 10 min at 4 °C. Finally, the protein was precipitated with cold 15% TCA for 2 h at −20 °C. After centrifugation at 4 °C for 10 min, the supernatant was discarded. The remaining precipitate was washed three times with cold acetone. The protein was redissolved in buffer (8 M urea, 100 mM TEAB, pH 8.0), and the protein concentration was determined with a 2-D Quant kit according to the manufacturer’s instructions.

### Trypsin Digestion and TMT Labeling

For digestion, the protein solution was reduced with 10 mM DTT for 1 h at 37 °C and alkylated with 20 mM IAA for 45 min at room temperature in dark. For trypsin digestion, the protein sample was diluted by adding 100 mM TEAB to reduce the urea concentration to less than 2 M. Finally, trypsin was added at a 1:50 trypsin-to-protein mass ratio for the first digestion, which was conducted overnight, and at a 1:100 trypsin-to-protein mass ratio for a second 4 h digestion. Approximately 100 μg of protein from each sample was digested with trypsin for the subsequent experiments. After trypsin digestion, the peptides were desalted using a Strata X C18 SPE column (Phenomenex) and vacuum-dried. The peptides were reconstituted in 0.5 M TEAB and processed according to the manufacturer’s protocol using a 6-plex TMT kit. Briefly, one unit of TMT reagent (defined as the amount of reagent required to label 100 μg of protein) was thawed and reconstituted in 24 μL of ACN. The peptide mixtures were then incubated for 2 h at room temperature and pooled, desalted and dried by vacuum centrifugation.

### HPLC Fractionation

The sample was fractionated by high-pH reverse-phase HPLC using an Agilent 300 Extend C18 column (5 μm particles, 4.6 mm ID, 250 mm in length). Briefly, the peptides were first separated with a gradient of 2% to 60% acetonitrile in 10 mM ammonium bicarbonate, pH 10, over 80 min, resulting in 80 fractions. The peptides were then combined into 18 fractions and dried by vacuum centrifugation.

### LC-MS/MS Analysis

The peptides were dissolved in 0.1% FA and then directly loaded onto a reversed phase pre-column (Acclaim PepMap 100, Thermo Scientific). Peptide separation was performed using a reversed-phase analytical column (Acclaim PepMap RSLC, Thermo Scientific). The gradient consisted of an increase from 6% to 22% of solvent B (0.1% FA in 98% ACN) over 19 min, then an increase from 22% to 35% over 13 min, followed by an increase to 80% over 4 min and finally, a hold at 80% for the last 4 min; the entire separation was performed at a constant flow rate on an EASY-nLC 1000 UPLC system. The resulting peptides were analysed by a Q ExactiveTM plus hybrid quadrupole-Orbitrap mass spectrometer (ThermoFisher Scientific). The peptides were subjected to an NSI source followed by tandem mass spectrometry (MS/MS) using a Q ExactiveTM plus instrument (Thermo) coupled online to the UPLC. The intact peptides were detected by the Orbitrap at a resolution of 70,000 and then selected for MS/MS using an NCE setting of 30; the ion fragments were detected by the Orbitrap at a resolution of 17,500. A data-dependent procedure that alternated between one MS scan followed by 20 MS/MS scans was applied for the top 20 precursor ions that were above an ion count threshold of 1E4 in the MS survey scan with 30.0 s of dynamic exclusion. The applied electrospray voltage was 2.0 kV. Automatic gain control (AGC) was used to prevent overfilling of the ion trap; 5E4 ions were accumulated for the generation of MS/MS spectra. For the MS scans, the m/z scan range was 350 to 1800. The fixed first mass was set to 100 m/z.

### Database Search

The resulting MS/MS data were processed using MaxQuant with an integrated Andromeda search engine (v.1.5.2.8). The tandem mass spectra were queried against the TRANS database, which was concatenated with a reverse decoy database, for *Magnolia officinalis*. Trypsin/P was specified as the cleavage enzyme, with allowance for up to 2 missing cleavages. The mass error was set to 10 ppm for precursor ions and 0.02 Da for fragment ions. Carbamidomethylation on Cys was specified as a fixed modification, and oxidation on Met and acetylation on protein N-terminals were specified as variable modifications. The false discovery rate (FDR) thresholds for proteins, peptides and modification sites were set to 1%. The minimum peptide length was set to 7. TMT-6plex was selected as the quantification method. All of the other parameters in MaxQuant were set to the default values.

### Validation of DEPs in Chlorophyll metabolism pathway of four cultivars by PRM

The protein expression levels obtained from the TMT analysis were confirmed by quantifying the expression levels of 7 selected proteins (Table 2) by a PRM-MS analysis carried out at the Jingjie PTM-Biolab Co., Ltd. (Hangzhou). The signature peptides for the target proteins were defined according to the TMT data, and only unique peptide sequences were selected for the PRM analysis. When there was no unique peptide in MPPMF, a repeat of the quantification was performed using PRM (Table 1). The proteins (60 μg) were prepared, reduced, alkylated, and digested with trypsin according to the protocol for TMT analysis. The obtained peptide mixtures were introduced into the mass spectrometer via a C18 trap column (0.10 × 20 mm; 3 μm) then a C18 column (0.15 × 120 mm; 1.9 μm). The MS measurements were performed using a quadrupole mass filter-equipped bench-top Orbitrap mass spectrometer (Q-Exactive; Thermo Scientific). The raw data were then analysed using Proteome Discoverer 1.4 (Thermo Fisher Scientific). The FDR was set to 0.01 for the proteins and peptides. Skyline 2.6 software was used for quantitative data processing and proteomic analysis.

**Table 1.**
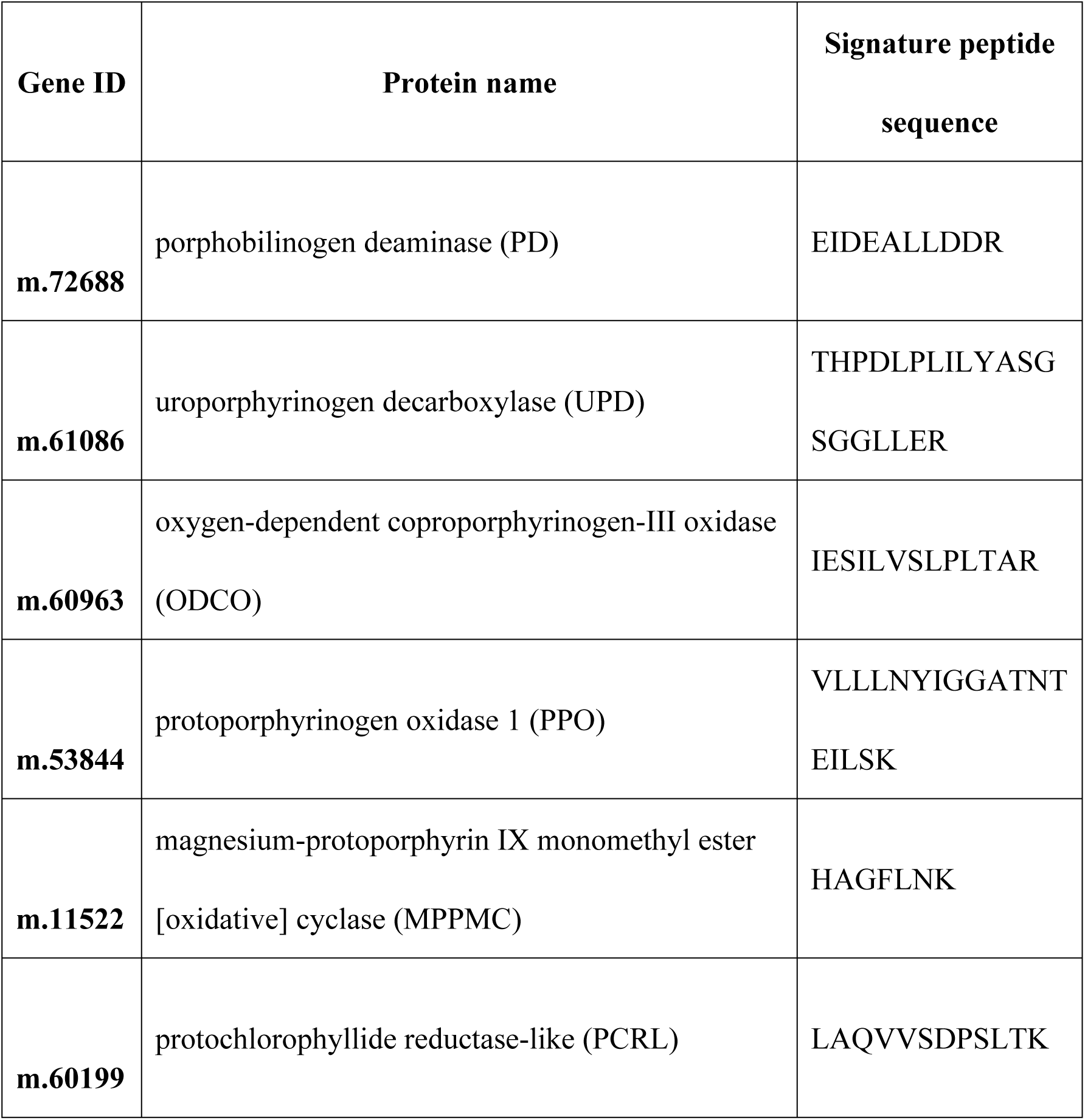
The unique peptides of PD, UPD, ODCO, PPO, MPPMC, RCRL

**Table 2.** Selected protein and peptides sequence used in research of parallel reaction monitoring

**Tabel 2** Selected protein and peptides sequence used in research of parallel reaction monitoring
| **Gene ID in transcriptome database** | **Protein name** | **Signature peptide sequence** |
| --- | --- | --- |
| **m.72688** | porphobilinogen deaminase (PD) | ILNQPLADIGGK |
| **m.61086** | uroporphyrinogen decarboxylase (UPD) | EFVPEESAPYVGEALT<br>ILR |
| **m.60963** | oxygen-dependent coproporphyrinogen-III oxidase (ODCO) | SVIPAYIPLIER |
| **m.53844** | protoporphyrinogen oxidase 1 (PPO) | LIEPFCSGVYAGDPSK |
| **m.58550** | magnesium protoporphyrin IX methyltransferase (MPPMF) | IGELFPGPSK |
| **m.11522** | magnesium-protoporphyrin IX monomethyl ester [oxidative] cyclase (MPPMC) | NLNQSEFEALLQEFK |
| **m.60199** | protochlorophyllide reductase-like (PCRL) | EHIPLFR |

## Results

### Leaf color results

DA, XA, LY and CH are the four main *Magnolia officinalis* cultivars in Sichuan province in China. During the developmental stages of their leaves, the colour has clear differences (Fig 1). The leaves of the DA cultivar are of a much deeper green than those of the other three cultivars. Subsequent measurements indicated that DA leaves have greater amounts of chlorophyll than those of XA, LY and CH (Table 3).

**Table 3.** The chlorophyll a and chlorophyll b content in four cultivars of *Magnolia officinalis*

| **Cultivar** | **Chlorophyll a**<br>**(mg/g FW)** | **Chlorophyll b**<br>**(mg/g FW)** | **Total chlorophyll**<br>**(mg/g FW)** |
| --- | --- | --- | --- |
| **CH** | 13.34±0.76 | 4.73±0.27 | 18.07±1.01 |
| **LY** | 18.09±0.13 | 7.17±0.56 | 25.26±0.59 |
| **XA** | 21.00±0.44 | 9.15±0.18 | 30.15±0.27 |
| **DA** | 26.36±0.22 | 11.21±0.17 | 37.56±0.40 |

### RNA-seq and proteomics results

Approximately 27 Gb of sequences in total were generated on the Illumina HiSeq sequencing platform during this study, which resulted in the assembly of 125,100 unigenes from all of the samples after filtering. The total length, average length, N50, and GC content of the unigenes were 110,568,181 bp, 883 bp, 1,660 bp and 43.58%, respectively. Moreover, 55,268 (NR: 44.18%), 39,362 (NT: 31.46%), 36,425 (Swissprot: 29.12%), 44,667 (KOG: 35.71%), 42,413 (KEGG: 33.90%), 18,597 (GO: 14.87%), and 46,330 (InterPro: 37.03%) unigenes were annotated by alignment with 7 functional databases. For the results of the functional annotation, 55,338 CDS were detected by Transdecoder. Finally, 37,588 SSRs distributed on 27,921 unigenes were detected, and 1,657 transcription factor sequences (TF) were predicted to be encoded in the unigenes (The raw data was supplied to NCBI, and the SRA accession ID: PRJNA492971). A total of 6,012 protein groups were identified, among which there were 5,063 quantified proteins that had been previously identified in a proteomic study. The fold-change cut-off was set so that proteins with quantitative ratios above 1.5 or below 0.67 were deemed significant. A total of 139 proteins were upregulated and 129 proteins were downregulated in group LY/CH. A total of 485 proteins were upregulated and 346 proteins were downregulated in group DA/CH. A total of 54 proteins were upregulated and 83 proteins were downregulated in group XA/CH. A total of 511 proteins were upregulated and 398 proteins were downregulated in group DA/LY. A total of 92 proteins were upregulated and 164 proteins were downregulated in group XA/LY. A total of 291 proteins were upregulated and 453 proteins were downregulated in group XA/DA(Supplementary file 1).

### Analysis of DEGs via comparison of the four cultivars

The comparison of four cultivars yielded the following results: 11,192 differentially expressed genes (DEGs) (5,142 upregulated and 6,050 downregulated) were found in CH-VS-DA, 11,962 DEGs (6,945 upregulated and 5,017 downregulated) were found in CH-VS-LY, 9,305 DEGs (6,348 upregulated and 2,957 downregulated) were found in CH-VS-XA, 14,315 DEGs (5,506 upregulated and 8,859 downregulated) were found in LY-VS-DA, 9,940 DEGs (5,644 upregulated and 4,296 downregulated) were found in LY-VS-XA, and 11,502 DEGs (3,478 upregulated and 8,024 downregulated) were found in XA-VS-DA (Fig 2).

### Important gene cross-talk between protein and transcription levels

#### Analysis of cross-talk between the whole transcriptome and the proteome

In this study, the cross-talk that was present between the proteome and the transcriptome was investigated. A total of 6,012 proteins were identified, among which 5,063 quantified proteins had been previously identified in a proteomic study. The fold-change cut-off was set so that proteins with quantitative ratios above 1.5 or below 0.67 were deemed significant. In the transcriptomic analysis, 96,664 transcripts were identified. Transcripts with a fold change above 1 and a corrected p value < 0.01 were deemed to be significantly differentially expressed. Correlation analysis of protein and transcript levels was performed, and the following results were obtained: 114 genes were downregulated and 182 genes were upregulated in both DA and CH, 170 genes were downregulated and 217 genes were upregulated in both DA and LY, and 154 genes were downregulated and 108 genes were upregulated in both DA and XA (Figs 3 and 4). Subsequently, analysis of the cross-talk between proteins and transcripts was performed for 113 genes involved in chlorophyll synthesis, with the following results: 1 gene was downregulated and 1 gene was upregulated for the comparison of DA and CH, 2 genes were downregulated and 9 genes were upregulated for the comparison of DA and LY, and 8 genes were downregulated and 1 gene was upregulated for the comparison of XA and DA (Figs 5 and 6); the detailed list of transcripts and proteins can be found in Supplementary files 2 and 3.

#### Functional enriched clustering of co-regulated (up-and-down regulated) protein categories

The function of co-regulated proteins or transcripts was analysed on a global level based on enriched clustering of GO and KEGG pathways (Figs 7-10). Interestingly, the proteins involved in tetrapyrrole- and porphyrin-containing compound biosynthetic pathways and in porphyrin and chlorophyll metabolism, all of which involve chlorophyll synthesis, were clearly different in XA, LY, and CH versus DA.

Specifically, chlorophyll-related proteins were selected that are involved in the same function or pathway. Among all of the identified proteins, 113 proteins involved in chlorophyll biosynthesis were found. For these 113 proteins, we performed further analysis of their biological processing, cellular components, molecular function, and participation in the KEGG pathway (Figs 11-14). Among these different proteins, those involved in porphyrin−containing compound metabolism, tetrapyrrole biosynthetic processing, porphyrin−containing compound biosynthetic processing, chlorophyll metabolism, the thylakoid, the photosynthetic membrane, porphyrin metabolism and photosynthesis in DA are clearly present at different levels than the corresponding proteins in XA, LY, and CH.

### Confirmation of chlorophyll metabolism pathway-related genes in four cultivars by PRM

According to analysis of cross-talk between proteins and transcripts, PD, UPD, ODCO, PPO, MPPMF, MPPMC, and RCRL have the highest expression levels among the four cultivars (Fig 15). These proteins are all rate-limiting enzymes involved in chlorophyll synthesis that have a direct effect on leaf colour. To confirm the PRM results, unique peptides derived from PD, UPD, ODCO, PPO, MPPMC, and RCRL (Table 1) were selected to be re-quantified by PRM. Because there was no unique peptide available, we did not re-quantify MPPMF. Our results verified that PD, UPD, ODCO, PPO, MPPMC, and RCRL have higher expression levels in DA than in the other three cultivars (Fig 16).

## Discussion

Nitrogen is an important component in the synthesis of chlorophyll and other photosynthetic proteins; approximately 75% of the nitrogen in plants is concentrated within chloroplasts, where it is mostly utilized in the construction of photosynthetic organs, making it a key factor in photosynthetic metabolism and plant growth [7,8]. The concentration of nitrogen per leaf mass is one of the most important leaf trait indicators and is closely related to the photosynthetic capacity of leaves and water use efficiency [9,10].

The results showed that the total amount of chlorophyll was 37.56 and 18.07 mg/g FW in DA and CH, respectively. The KEGG pathway analysis showed that some key genes involved in porphyrin, chlorophyll and nitrogen metabolism were upregulated. These genes included those encoding glutamyl-tRNA reductase, protochlorophyllide reductase, magnesium-chelatase subunit ChlH, chloroplastic magnesium-protoporphyrin IX monomethyl ester [oxidative] cyclase, glutamate synthase, and ferredoxin-nitrite reductase. Glutamyl-tRNA reductase (GluTR) catalyses the NADPH-dependent reduction of Glu-tRNA^Glu^ to GSA, which plays a vital role in chlorophyll biosynthesis [11]. Moreover, the results showed that the chlorophyll content in DA was much higher than that in CH. The upregulation of GluTR was consistent with the amount of chlorophyll that was measured.

Nitrogen, as one of the components of enzymes, amino acids, nucleic acids, and chlorophyll, plays an essential role in plant growth and development [12]. Chlorophyll (Chl) is the most important photosynthetic pigment in the chloroplast and plays an essential role in absorbing light and transferring light energy within the antenna systems [13]. Chlorophylls a and b are important components of chloroplast pigments in higher plants, accounting for approximately 75% of total chloroplast pigment [14]. The effects of nitrogen on plant growth and development are very obvious. When nitrogen is sufficient, plants can synthesize more proteins and promote cell division and growth. In this study, the results showed that ferredoxin-nitrite reductase was upregulated in DA compared with CH. The differences in chlorophyll content also supported our results. Moreover, the results showed that some key genes involved in porphyrin, chlorophyll and nitrogen metabolism were upregulated in DA compared with LY. These genes included those encoding magnesium-chelatase subunit ChlI, glutamyl-tRNA reductase, protochlorophyllide reductase, magnesium-protoporphyrin IX monomethyl ester [oxidative] cyclase, magnesium-chelatase subunit ChlH, geranylgeranyl diphosphate reductase, ferredoxin--nitrite reductase, and glutamine synthetase leaf isozyme. The upregulated genes were consistent with the chlorophyll contents in DA and LY, respectively.

Auxins, which comprise a class of plant hormones, influence cellular expansion and differentiation and ethylene biosynthesis, and thereby affect nearly all aspects of plant growth and development [15,16]. Bajguz A (2014) showed that 1μM NAA (auxin) induced an increase in cell density of 24% after 72 h of growth in *Chlorella vulgaris* [17]. In the present study, indole-3-acetaldehyde oxidase, which is involved in auxin biosynthesis, had a pattern of continuously upregulated expression in XA; the chlorophyll amount was consistent with this pattern. Bioactive gibberellins (GAs) play essential roles in regulating a variety of developmental processes in plants, including seed germination, leaf senescence, vegetative growth, and fruit development [18,19]. Moreover, it has been reported that GAs stimulate plant growth by promoting cell elongation in plants [20]. The gibberellin receptor GID1C, which is involved in the auxin signal pathway, had a continuously upregulated expression pattern in DA. The relative quantification of the protein values was 0.984, 1, 0.978 and 1.111 in LY, CH, XA and DA, respectively. The upregulation of the gibberellin receptor may be attributed to its ability to increase the levels of chlorophyll. ABA plays a positive role in the process of leaf senescence, and exogenously-applied ABA has been shown to accelerate chlorophyll degradation in Arabidopsis [21]. Moreover, it has long been known that exogenously applied ABA can induce the expression of senescence-associated mRNAs and lead to decreases in chlorophyll content in detached leaves [22]. According to the current study, the abscisic acid receptor PYL8, which is involved in the ABA signalling pathway, had a continuously downregulated expression pattern in DA. The values of the relative quantification of the protein were 1.06, 0.818, 0.891 and 0.976 in LY, CH, XA and DA, respectively.

With the data provided by transcriptomic and proteomic profiling, a number of important genes involved in the development and regulation of chlorophyll metabolism were analysed, which can further contribute to the understanding of colour changes in the four cultivars of *Magnolia officinalis*. Importantly, some of the genes involved in the metabolism of plant hormone metabolism also play important roles in chlorophyll metabolism; the up- or down-regulation of these genes or proteins may be attributed to their ability to increase the levels of chlorophyll.

## Conclusions

In summary, proteomics research not only provides new insights into protein expression patterns but also enables the identification of many attractive candidates for further research into the effects of pesticides on chlorophyll metabolism in *Magnolia officinalis*. Based on this work, the phenomenon behind the greater amount of green colour in leaves of the DA cultivar is clear. The transcriptome and proteome analysis of the four main *Magnolia officinalis* cultivars in Sichuan province in China identified 125,100 unigenes and 6012 proteins. According to the analysis of cross-talk between the transcriptome and the proteome, proteins involved in chlorophyll metabolism (listed as Table 2) have a higher expression level in DA than in the other three cultivars. Subsequently, PRM analysis confirmed that the expression level of these proteins is greater in DA than in XA, LY, and CH.

## Supporting information

**Fig. 1** Leaves color’s difference of CH, LY, XA, DA.

**Fig. 2** The summary of DEGs in comparable group between DA, LY, CH, XA.

**Fig. 3** the relationship between the whole transcriptome and proteome.

**Fig. 4** Significance differentially expressed proteins or transcriptions compare bar diagram.

**Fig. 5** the relationship between transcriptome and proteome involved chlorophyll mechanism.

**Fig. 6** Significance differentially expressed proteins or transcriptions involved chlorophyll machanism compare bar diagram.

**Fig. 7** Biological analysis of genes co-regulated on transcriptome and proteome.

**Fig. 8** Cellular compotent analysis of genes co-regulated on transcriptome and proteome.

**Fig. 9** Molecular function analysis of genes co-regulated on transcriptome and proteome.

**Fig. 10** KEGG pathway analysis of genes co-regulated on transcriptome and proteome.

**Fig. 11** Biological analysis of genes co-regulated involved chlorophyll mechanism on transcriptome and proteome.

**Fig. 12** Cellular compotent analysis of genes co-regulated involved chlorophyll mechanism on transcriptome and proteome.

**Fig. 13** Molecular function analysis of genes involved chlorophyll mechanism co-regulated on transcriptome and proteome.

**Fig. 14** KEGG pathway analysis of genes co-regulated involved chlorophyll mechanism on transcriptome and proteome.

**Fig. 15** Quantification and identification of proteins by PRM. **a)** The re-quantification and re-identification result of protein PD based on peptides sequence ILNQPLADIGGK; **b)** The re-quantification and re-identification result of protein UPD based on peptides sequence EFVPEESAPYVGEALTILR; **c)** The re-quantification and re-identification result of protein ODCO based on peptides sequence SVIPAYIPLIER; **d)** The re-quantification and re-identification result of protein PPO based on peptides sequence LIEPFCSGVYAGDPSK; **e)** The re-quantification and re-identification result of protein MPPMF based on peptides sequence IGELFPGPSK; **f)** The re-quantification and re-identification result of protein MPPMC based on peptides sequence NLNQSEFEALLQEFK; **g)** The re-quantification and re-identification result of protein MPPMC based on peptides sequence EHIPLFR.

**Fig. 16 a)** The re-quantification and re-identification result of protein PD based on peptides sequence ILNQPLADIGGK; **b)** The re-quantification and re-identification result of protein UPD based on peptides sequence EFVPEESAPYVGEALTILR; **c)** The re-quantification and re-identification result of protein ODCO based on peptides sequence SVIPAYIPLIER; **d)** The re-quantification and re-identification result of protein PPO based on peptides sequence LIEPFCSGVYAGDPSK; **e)** The re-quantification and re-identification result of protein MPPMC based on peptides sequence NLNQSEFEALLQEFK; **f)** The re-quantification and re-identification result of protein RCRL based on peptides sequence EHIPLFR.

## Author Contributions

**Conceptualization:**Jin Pei.

**Data curation:**Jiang Chen.

**Formal analysis:**Qinghua Wu, Jin Pei.

**Funding acquisition:** Qinghua Wu, Jin Pei.

**Investigation:** Qinghua Wu, Dan Wei, Yuping Liu, Qianqian Liu.

**Methodology:** Qinghua Wu, Jiang Chen.

**Project administration:** Qinghua Wu, Jin Pei.

**Resources:** Qinghua Wu, Dan Wei.

**Software:** Jiang Chen, Cuiping Chen, Chaoxiang Ren.

**Supervision:** Jin Pei.

**Validation:** Qinghua Wu, Dan Wei, Yuping Liu, Qianqian Liu.

**Visualization:** Jiang Chen, Chaoxiang Ren.

**Writing – original draft:** Qinghua Wu, Dan Wei.

**Writing – review & editing:** Qinghua Wu, Dan Wei, Yuping Liu, Qianqian Liu, Cuiping Chen, Chaoxiang Ren, Jiang Chen, Jin Pei.

**Competing interests:**The authors declare that they have no competing interests.

